# Fitness effects of new mutations in *Chlamydomonas reinhardtii* across two stress gradients

**DOI:** 10.1101/033886

**Authors:** Susanne A. Kraemer, Andrew D. Morgan, Robert W. Ness, Peter D. Keightley, Nick Colegrave

## Abstract

Most spontaneous mutations affecting fitness are likely to be deleterious, but the strength of selection acting on them might be impacted by environmental stress. Such stress-dependent selection could expose hidden genetic variation, which in turn might increase the adaptive potential of stressed populations. On the other hand, this variation might represent a genetic load and thus lead to population extinction under stress. Previous studies to determine the link between stress and mutational effects on fitness, however, have produced inconsistent results. Here, we determined the net change in fitness in 29 genotypes of the green algae *Chlamydomonas reinhardtii* that accumulated mutations in the near absence of selection for approximately 1,000 generations across two stress gradients, increasing NaCl and decreasing phosphate. We found mutational effects to be magnified under extremely stressful conditions, but such effects were specific both to the type of stress as well as to the genetic background. The detection of stress-dependent fitness effects of mutations depended on accurately scaling relative fitness measures by generation times, thus offering an explanation for the inconsistencies among previous studies.

## Introduction

Spontaneous mutation is a fundamental biological process that generates the abundant variation we observe in nature. Understanding the fitness effects of new mutations is central, therefore, to many problems in evolutionary biology. Whilst knowledge of both the rate and fitness effects of spontaneous mutation has increased for several species (Drake et al., 1998; Lynch et al., 1999; Eyre-Walker and Keightley, 2007), our understanding of the extent by which mutational effects depend on the environment is limited. Environment-specific fitness effects of genotypes (GxE interactions) are commonly observed for standing genetic variation in natural populations (Kassen and Bell, 2000; Kraemer and Kassen, 2015). Such GxE interactions may be caused by recently arisen mutations that have environment-specific fitness effects. Alternatively, GxE interactions may be actively maintained by selection, for example in the form of local adaptation to different environments (Belotte et al., 2003). Tests for selection acting on standing genetic variation in natural populations are usually difficult to perform and thus the degree to which GxE interactions are adaptive is largely unknown. A related idea is whether stress might release cryptic genetic variation, which selection can subsequently act upon (Latta et al., 2015). Such a release of variation might, for example, be caused by mutations with effects that are too small to be selected under benign conditions, but effectively selected under stressful conditions (Lenski and Remold, 2001; Kishony and Leibler, 2003). This phenomenon may manifest itself as an increase in the expressed genetic variation among genotypes when tested in a stressful environment compared to a more benign environmental control.

It is widely believed that the vast majority of new mutations that affect fitness are deleterious (Keightley and Lynch, 2003; Loewe and Hill, 2010, but see Rutter et al., 2010) and it has often been suggested that the deleterious effects of new mutations may be amplified in stressful environments (Agrawal and Whitlock, 2010; Kondrashov and Houle, 1994; Korona, 1999; Martin and Lenormand, 2006). If this is the case, inference of the properties of new mutations, typically made in relatively benign laboratory conditions, may provide a misleading view of their fitness effects in natural populations, where conditions are generally assumed to be more stressful. There is some empirical support for the existence of stronger fitness effects of spontaneous mutations in stressful environments, with examples in *E. coli* (Remold and Lenski, 2001; Cooper et al., 2005), *S. cerevisae* (Szafraniec et al., 2001; Jasnos et al., 2008), *Drosophila* (Kondrashov and Houle, 1994; Fry and Heinsohn, 2002; Wang et al., 2009; Young et al., 2009), *C. elegans* (Vassilieva et al., 2000) and *C. briggsae* (Baer et al., 2006). However, the generality of such observations has been brought into question by a number of studies, particularly in microbial systems (Korona, 1999; Kishony and Leibler, 2003; Jasnos et al., 2008;), showing that some forms of environmental stress can actually reduce the effects of deleterious mutations (reviewed in Agrawal and Whitlock, 2010).

Previous studies of the phenotypic effects of mutations under stressful conditions have typically examined the performance of genotypes in pairs of environments that are categorically different, for example environments with or without a antibiotic (Kishony and Leibler, 2003), or high *versus* low temperature (Szafraniec et al., 2001). The genetic basis of performance in such environments may differ dramatically in ways that are unrelated to the severity of the stress, and this may explain some of the inconsistency seen in previous studies. For example, suppose that there are large differences in the number of genes expressed in different environments, but that the number of genes expressed is uncorrelated with stress. If we compare performance in arbitrary pairs of environments, then any differences in performance may simply reflect differences in the size of the mutational target for fitness in these environments, rather than effects of stress *per se*. Furthermore, extreme laboratory environments are unlikely to represent conditions generally encountered by natural populations. Rather, changing conditions are likely to occur due to environmental deterioration across a gradient, stretching from benign, to slightly stressful, to detrimental and finally lethal. Moreover, different stressors might elicit different responses, depending on which aspect of the organism’s physiology they act on.

Here, we utilize lines derived in the absence of (most) selection during the course of a mutation accumulation (MA) experiment (Morgan et al., 2014). Specifically, we study 29 spontaneous MA lines derived from two genetically distinct strains of the unicellular green algae *Chlamydomonas reinhardtii*, along with the ancestral strains they were derived from, to investigate change in the fitness effects of *de novo* mutations as two environmental stressors are increased along a continuum. *Chlamydomonas reinhardtii* shares characteristics with both microbial as well as multicellular species. For example, its generation time and effective population size are close to those of *E. coli* and *S. cerevisiae* (Ness et al., 2012), whereas its genome size and structure are comparable to multicellular eukaryotes. In this study, we use genotypes generated by ca. 1,000 generations of mutation accumulation to address the following questions: 1) Are mutational effects ameliorated or exaggerated under stressful conditions? 2) Can we observe GxE interactions in genotypes derived from a mutation accumulation regime? 3) Does increasing stress lead to a release of cryptic variation?

We found that severe stress can amplify the effects of new mutations, but such condition dependence was impacted both by the type of stress as well as the genetic background. The detection of such subtle effects depended on accurate scaling of fitness by ancestral generation times, thus providing a partial explanation for the inconsistency among results from previous studies.

## Materials and Methods

### Mutation accumulation lines and ancestral strains

We studied MA lines and their ancestors derived from two genetically distinct laboratory strains of *C. reinhardtii*. Full details of the production of the MA lines are provided elsewhere (Morgan et al., 2014). There were 15 and 14 MA lines from ancestral strains CC-2344 and CC-2931, respectively. Briefly, the MA lines were generated under conditions where the effectiveness of natural selection is reduced by bottlenecking to a single cell at regular intervals (approximately every 12 divisions) for ~1,000 generations. Full genome sequencing showed that the MA lines have accumulated substantial numbers of new mutations (Ness et al. 2015). They typically show reduced growth rate compared to their ancestors when assayed in a benign laboratory environment (Morgan et al., 2014). The ancestors and their descendant MA lines had all been cryopreserved in liquid nitrogen prior to this study.

### Reconditioning of MA lines prior to the fitness assays

Prior to the fitness assays, we allowed cryopreserved cultures of all strains to thaw in a water bath at 35^o^C for 5 minutes. We transferred the cultures to 30ml universal tubes containing 10ml of Bold’s liquid media (Bold, 1942) and incubated them in standard growth conditions (25^o^C under constant white light illumination) for 7 days. We then transferred 100μl of each MA line culture into a new universal tube containing 5ml of Bold’s media. At the same time, we created replicates of the two ancestors by inoculating 15 samples of each into separate tubes of media. These cultures were incubated under standard growth conditions for four days. Finally, we transferred 2μl of each culture to individual wells containing 200μl of Bold’s media on a 96 well plate. Each MA line was randomly paired with one of the replicate cultures of the corresponding ancestor, and these two cultures were allocated at random to adjacent wells. In order to avoid edge effects (Morgan et al., 2014), we only used the central 60 wells of each plate, and filled the outer wells with Bold’s media to maintain humidity and to reduce evaporation from the central wells. We repeated this procedure three times, to create three independent conditioning plates with a different random allocation of genotypes on each plate. Subsequently, we incubated these plates for three days prior to the fitness assays.

### Manipulating environmental stress

We created two environmental stress gradients by modifying the concentrations of components of standard Bold’s media that are known to affect the performance of *C. reinhardtii*: NaCl and phosphate (Bell, 1992; Goho and Bell, 2000; Moser and Bell, 2011). For the phosphate gradient, we altered the concentration of the dipotassium phosphate (K_2_HPO_4_) and the monopotassium phosphate (KH_2_PO_4_) components so that the media contained 400% (0.6 g/L), 200% (0.3 g/L), 100% (0.15 g/L), 50% (0.075 g/L), 25% (0.038 g/L), 10% (0.015 g/L), and 5% (0.0075 g/L) of the concentration of phosphate of standard Bold’s media (0.15 g/L). For the NaCl gradient, we supplemented standard liquid Bold’s medium with between 0 and 6gl^−1^ of NaCl in 1gl^−1^ increments. Higher concentrations of NaCl investigated in pilot studies largely failed to support growth. Natural genotypes of *C. reinhardtii* vary in their performance over these ranges, and growth is significantly reduced in the most stressful environments we investigated (the lowest phosphate or the highest NaCl).

Before measuring the fitness of the lines in the different assay conditions we first allowed them to acclimatize in order to ensure that we measured fitness, rather than differences in plasticity in response to a new environment. We set up three replicate 96 well plates per environmental condition (i.e., level of phosphate or NaCl) by adding 200μl of liquid media. We initiated the fitness assays by adding 2μl of each culture from one conditioning plate. A different conditioning plate was used to initiate each replicate plate for a given environment. These plates were incubated under standard conditions for four days to allow the lines to acclimatize to the assay conditions.

After acclimatizing the lines, we estimated the maximum growth rates of the MA lines and their ancestors in 96 well plates containing the appropriate assay media. 2μl of each culture was then transferred to the corresponding well of a second 96 well plate containing the same growth media. This maintained the randomization of the lines across the 96 well plates and pairing with their ancestors for the assay. These plates were incubated for four further days under standard growth conditions, and the absorbance of each well at 650nm was measured three times each day to provide estimates of culture density. Absorbance values below 0.01 do not provide reliable estimates of cell density, and this places a lower threshold on the culture density that we can estimate using this method. However, this threshold corresponds to a cell density of about 1 × 10^5^ cells per ml, which is well below the cell density at which exponential growth ceases (approximately 1 × 10^7^ cells per ml), and so this method allows us to estimate the maximum growth rate of cultures once they pass this detection threshold.

The maximum growth rate of each culture was calculated as

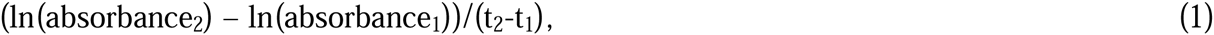
where t_1_ and t_2_ are the times in hours of the first two assays, at which an absorbance above 0.01 detection threshold of the plate reader was measured for a culture, which corresponds to the period of fastest growth under these assay conditions. absorbance_1_ and absorbance_2_ are the absorbance values at these two time point, respectively. Since cultures in different environmental conditions differed in their time to reach the absorbance threshold, the actual values of t_1_ and t_2_ differed for different cultures. Cultures that failed to grow were assigned a growth rate of 0.

For logistical reasons, it was not possible to assay both stress gradients concurrently. Instead, we first assayed all MA lines across the NaCl gradient, and carried out the assays for the phosphate gradient two weeks later.

### Statistical analyses

Data sets were analyzed separately for each stressor. To examine effects of mutation accumulation and environmental stress on growth rate, we fitted linear mixed models using the packages lme4 (Magezi, 2015) nlme in R (R Development Core Team, 2009). *p*-values were assigned using the Satterthwaite-approximation, based on SAS Proc mixed theory (Schaalje et al., 2002), as implemented in the lmerTest package in R.

Specifically, we investigated the relationship between relative fitness of MA genotypes and the level of environmental stress by fitting linear mixed models accounting for random effects due to plates and ancestry. We based calculations of relative fitness on the average growth of all pseudoreplicates of the ancestor per plate rather than using the individual values of each paired ancestor, because some ancestral samples failed to grow. All model coefficients can be found in the supplemental methods file.

Genotype-by-environment interactions for all genotypes between each possible pair of test environments were further investigated by partitioning GxE into two components: the genetic correlation of growth and the variation of the genetic standard deviation expressed by genotypes between the two environments (Robertson, 1959; Kassen and Bell, 2000; Kraemer and Kassen, 2015; Barrett et al., 2005). The genetic correlation of growth describes the GxE component that determines the consistency of performance of genotypes across environments. For example, if test environments are highly similar, the rank order positions of fitness of genotypes in the two environments are expected to be similar, leading to a high genetic correlation. Very dissimilar environments, on the other hand, are likely to elicit different fitness responses such that the rank order of fitness of genotypes changes substantially from one environment to another, leading to a low genetic correlation of growth. The second component of GxE interactions describes how the variance in performance among genotypes changes between the two environments and will be generated if more variation is expressed in one environment than the other.

Total GxE interactions are described by

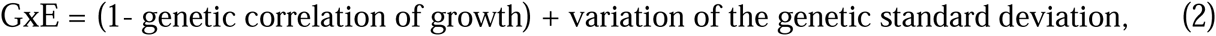
and formally by the Robertson equation (Robertson, 1959):

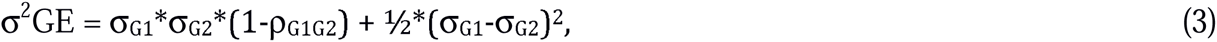
where σ_G1_ and σ_G2_ are the genetic standard deviations of the fitness of all genotypes grown in environment 1 and 2 specifically and *ρ*_G1G2_ is the genetic correlation of growth between the two environments. Thus, overall GxE, as well as the variation of the genetic standard deviation, are expected to increase with increasing environmental dissimilarity, while the genetic correlation of growth is expected to decrease. A schematic showing GxE variation due to either the genetic correlation of growth or due to the variation of the genetic standard deviation is shown in Supp. Fig. 1.

Since GxE measures in these correlations are the results of comparisons of pairs of environments and thus not necessarily independent data points, we assessed the significance of GxE patterns across environments using a permutation test in which, instead of assuming a normal distribution of residuals, we generated a distribution based on the data. Test results were subsequently compared to this distribution to assess significance. Specifically, we generated GxE component estimates for each pair of environments based on the Robertson equation with the asreml package in R and regressed these on environmental difference (the difference in either NaCl or Phosphate concentration between the two focal environments). We then generated 10,000 permuted data sets by randomly assigning the environmental difference scores to the pairwise component estimates without replacement and repeated the regression analysis on each. We subsequently utilized the resulting distribution regression statistics as a null-hypothesis to test for significant effects of total GxE, genetic correlation of growth and variation of the genetic standard deviation with respect to differences between environments

The expressed genetic variation was calculated based on the relative fitness values of MA genotypes under each stress level using the lme4 package in R (Magezi, 2015). Confidence intervals around values of expressed genetic variation were calculated assuming a chi-squared distribution.

## Results

### 1. Overall effect of the environments on growth rate

The growth rates of the MA lines and their ancestors were significantly reduced by decreasing phosphate and increasing NaCl concentrations (linear mixed models, effect of stress treatment on growth rate, all *p* < 0.001, Table 1, Figure 1). Across the gradient of decreasing phosphate concentration, growth was reduced by 65%, while increasing NaCl concentration reduced growth by 84%. Although the MA lines grew on average 20% more slowly than their ancestors across all conditions, we found no evidence that the stress treatments impacted MA line growth rates differently compared to paired ancestral pseudoreplicates (linear mixed models, interaction between type (MA or ancestor) and stress level, all *p* > 0.05, Table 1, Figure 1). Thus, the magnitude of the average reduction in maximum growth rate due to accumulated spontaneous mutations appears to be unaffected by stress.

**Table 1:**
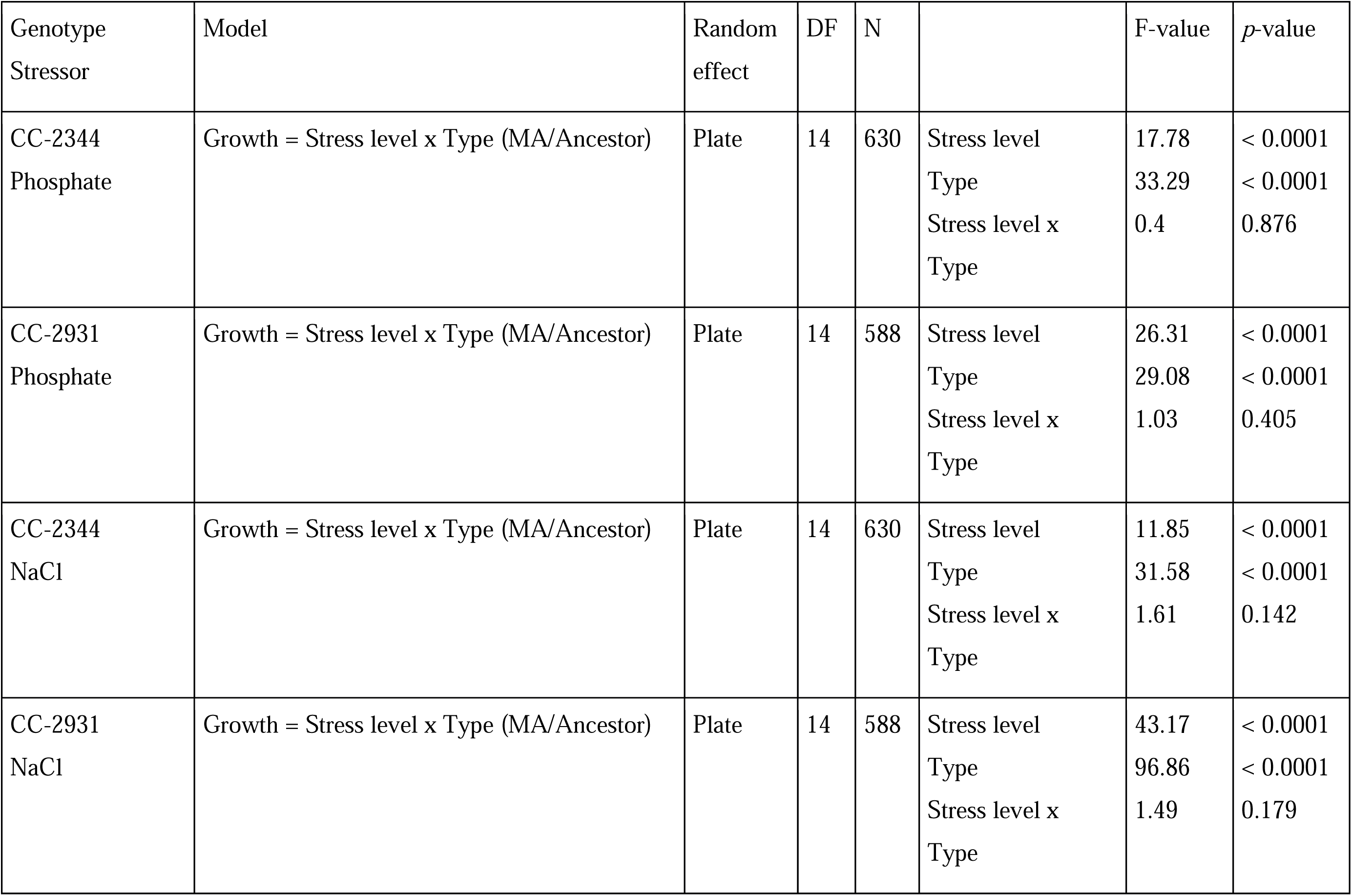
Linear mixed model analyses of maximum growth rate.

| Genotype<br>Stressor | Model | Random<br>effect | DF | N |  | F-value | *p*-value |
| --- | --- | --- | --- | --- | --- | --- | --- |
| CC-2344<br>Phosphate | Growth = Stress level x Type (MA/Ancestor) | Plate | 14 | 630 | Stress level<br>Type<br>Stress level x<br>Type | 17.78<br>33.29<br>0.4 | < 0.0001<br>< 0.0001<br>0.876 |
| CC-2931<br>Phosphate | Growth = Stress level x Type (MA/Ancestor) | Plate | 14 | 588 | Stress level<br>Type<br>Stress level x<br>Type | 26.31<br>29.08<br>1.03 | < 0.0001<br>< 0.0001<br>0.405 |
| CC-2344<br>NaCl | Growth = Stress level x Type (MA/Ancestor) | Plate | 14 | 630 | Stress level<br>Type<br>Stress level x<br>Type | 11.85<br>31.58<br>1.61 | < 0.0001<br>< 0.0001<br>0.142 |
| CC-2931<br>NaCl | Growth = Stress level x Type (MA/Ancestor) | Plate | 14 | 588 | Stress level<br>Type<br>Stress level x<br>Type | 43.17<br>96.86<br>1.49 | < 0.0001<br>< 0.0001<br>0.179 |

**Figure 1:**
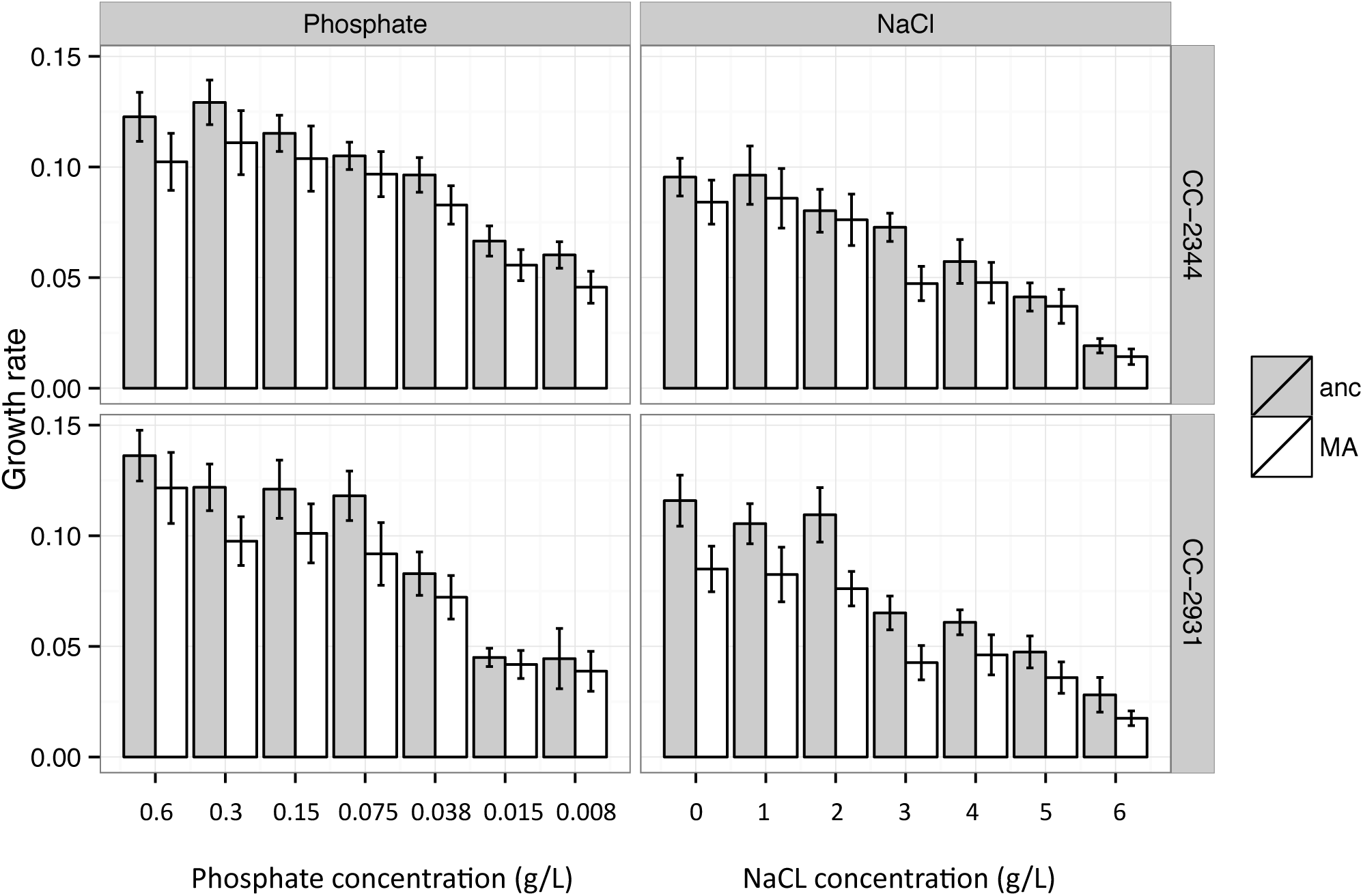
Mean growth rates (h^−1^) of the MA line genotypes and their respective ancestors under seven levels of two different stress regimes (400% (0.6gl^−1^), 200% (0.3gl^−1^), 100% (0.15gl^−1^), 50% (0.075gl^−1^), 25% (0.038gl^−1^), 10% (0.015gl^−1^), and 5% (0.0075gl^−1^) KH_2_PO_4_; and 0 to 6gl^−1^ of NaCl in 1gl^−1^ increments, ordered by increase in stress level. Error bars show 95% confidence intervals.

### 2. Does environmental stress affect relative fitness?

The evolutionary dynamics of a new mutation is not determined by its effect on growth rate *per se*, but instead by its effect on relative fitness compared to its ancestral genotype under the same environmental conditions (Chevin, 2011). Here, we focus on the growth rate of MA lines, relative to the growth rate of their ancestors, as a proxy for relative fitness (1 - selection coefficient [*s*]) of each MA line genotype under a given stress level. For a specific MA line genotype, we first calculated Δ*G* = its growth rate - average growth rate of all its ancestral pseudoreplicates on the same plate. Since both growth rates are measured on a log scale, this provides a measure of relative fitness in units of time. However, in evolutionary biology we are generally interested in relative fitness per generation. This distinction becomes particularly important in studies such as this where fitness is compared among environments where the generation time varies. To obtain a measure of relative fitness per generation (*s*), we then scaled Δ*G* by the ancestral generation time, measured under the same conditions, by multiplying it with ln(2) divided by the average ancestral growth rate on the same plate (Chevin, 2011). This measure of relative fitness should be directly comparable across our selection environments as well as providing estimates that can be compared to other organisms with different generation times. In contrast to results obtained using raw growth rates, we found that stress exacerbated the effects of mutations: we detected a weak but significant effect of increasing NaCl stress on relative fitness (linear mixed model, effect of stress level on relative fitness of all MA genotypes nested within their respective ancestor, *p* < 0.01, Table 2, Fig. 2).

**Table 2:** Linear mixed model analyses of relative fitness of all MA line genotypes under either phosphate or NaCl stress.

| Data set | Model | Random effects | DF | N |  | F-values | *p*-value |
| --- | --- | --- | --- | --- | --- | --- | --- |
| All<br>Phosphate | Relative fitness = Stress level x<br>Genotype | Plate<br>Ancestor | 104 | 609 | Stress level | 0.91 | 0.487 |
|  |  |  |  |  | Genotype | 2.98 | < 0.001 |
|  |  |  |  |  | Stress level x Genotype | 0.78 | 0.923 |
| All<br>NaCl | Relative fitness = Stress level x<br>Genotype | Plate<br>Ancestor | 104 | 609 | Stress level | 5.21 | 0.005 |
|  |  |  |  |  | Genotype | 6.89 | < 0.001 |
|  |  |  |  |  | Stress level x Genotype | 1.41 | 0.016 |

**Figure 2:**
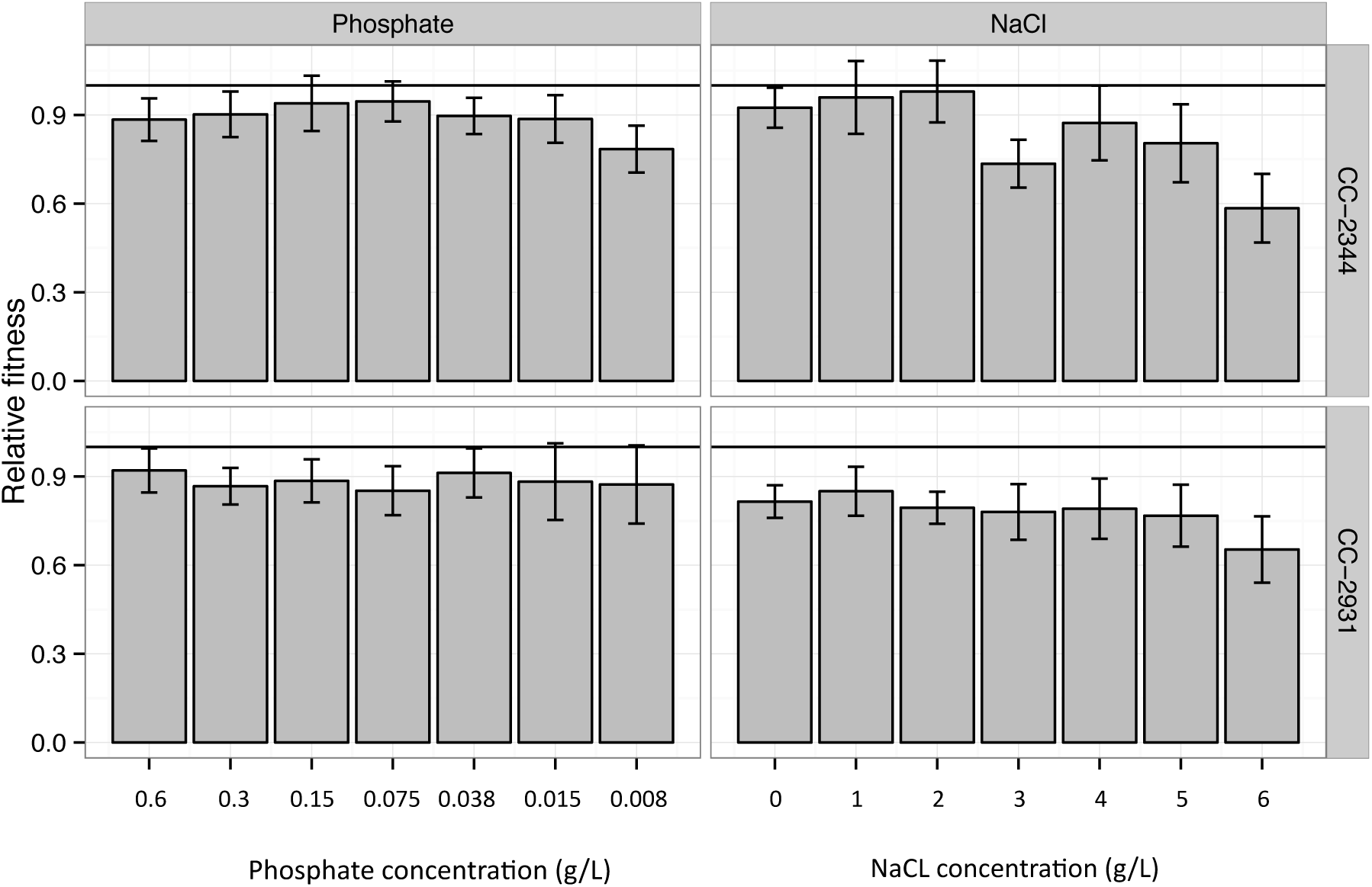
Mean relative fitness of MA line genotypes derived from two ancestral strains under decreasing phosphate 400% (0.6gl^−1^), 200% (0.3gl^−1^), 100% (0.15gl^−1^), 50% (0.075gl^−1^), 25% (0.038gl^−1^), 10% (0.015gl^−1^), and 5% (0.0075gl^−1^) KH_2_PO_4_; and increasing NaCl (0 to 6gl^−1^ of NaCl in 1gl^−1^ increments) conditions, ordered by increase in stress level. Error bars indicate 95% confidence intervals.

Although MA line genotypes derived from one of the ancestors, CC-2344, also showed a slight decrease in relative fitness under the highest phosphate stress, we did not detect a significant effect of stress level on relative fitness under this stress regime (linear mixed model, effect of stress level on relative fitness of all MA genotypes derived from both ancestors, all *p* = 0.15 and *p* = 0.55 respectively, Table 2, Fig. 2).

MA genotypes under NaCl stress showed a significant interaction between stress levels and genotype (Table 2, *p* < 0.05), indicating that MA lines respond differently to NaCl stress. We further investigated this GxE interaction between the individual MA genotypes and stress level by separately analyzing data from the MA line genotypes derived from the two ancestors. Only MA genotypes derived from ancestor CC-2344 assayed under NaCl stress exhibited a significant genotype-by-environment interaction (linear mixed models, interaction of stress level with MA line genotype, *p* < 0.05, all other interactions *p* > 0.05, Table 3). However, we did not detect any such effects for the two MA lines under phosphate stress (linear mixed models, interaction of stress level with each MA line genotype, *p* > 0.05, Table 3). We also performed all analyses based on relative fitness unscaled by ancestral generation time (1-*s*). Importantly, we did not detect any effect of stress level on fitness, nor any significant GxE interactions between genotype and stress level (Supp. Fig. 2, all *p* > 0.05).

**Table 3:** Linear mixed model analyses of relative fitness for data sets partitioned by stressor and ancestor.

| Data set | Model | Random effect | DF | N |  | F-value | *p*-value |
| --- | --- | --- | --- | --- | --- | --- | --- |
| 2344<br>Phosphate | Relative fitness =<br>Stress level x Genotype | Plate | 14 | 315 | Stress level | 1.11 | 0.403 |
|  |  |  |  |  | Genotype | 4.14 | < 0.001 |
|  |  |  |  |  | Stress level x Genotype | 1.21 | 0.147 |
| 2931<br>Phosphate | Relative fitness =<br>Stress level x Genotype | Plate | 14 | 294 | Stress level | 0.28 | 0.936 |
|  |  |  |  |  | Genotype | 2.18 | 0.012 |
|  |  |  |  |  | Stress level x Genotype | 0.97 | 0.547 |
| 2344<br>NaCl | Relative fitness =<br>Stress level x Genotype | Plate | 14 | 315 | Stress level | 6.71 | 0.002 |
|  |  |  |  |  | Genotype | 4.64 | < 0.001 |
|  |  |  |  |  | Stress level x Genotype | 1.60 | 0.004 |
| 2931<br>NaCl | Relative fitness =<br>Stress level x Genotype | Plate | 14 | 294 | Stress level | 0.76 | 0.614 |
|  |  |  |  |  | Genotype | 6.91 | < 0.001 |
|  |  |  |  |  | Stress level x Genotype | 0.92 | 0.656 |

### 3. How do GxE interactions change between environments with different stress levels?

To investigate how GxE interactions vary between environments, they can be partitioned into their components, genetic correlation of growth and the variation of the genetic standard deviation, according to the Robertson equation (Robertson, 1959). To investigate how GxE might be correlated with increasing stress levels, these components can be plotted against measures of environmental dissimilarity between each pair of environments tested, for example against the difference in NaCl or phosphate concentrations. Overall GxE interactions and the variation of the genetic standard deviations are expected to increase with increasing differences between environments, while the genetic correlation of growth, or how consistent MA genotypes behave across environments, is expected to decrease. Here, we focus on MA genotypes derived from ancestor CC-2344 under NaCl stress, which showed significant GxE interactions (Table 3, Fig. 3). We do not see a clear pattern with increasing differences between environments for neither total GxE interactions nor the genetic correlation of growth. The variation of the genetic standard deviation, however, increases significantly with increasing environmental differences (Permutation tests (N= 10,000) of GxE (*p* > 0.05), genetic correlation of growth (*p* > 0.05) and variation of the genetic standard deviation (*p* < 0.01) against differences in NaCl concentrations, Fig. 3).

**Figure 3:**
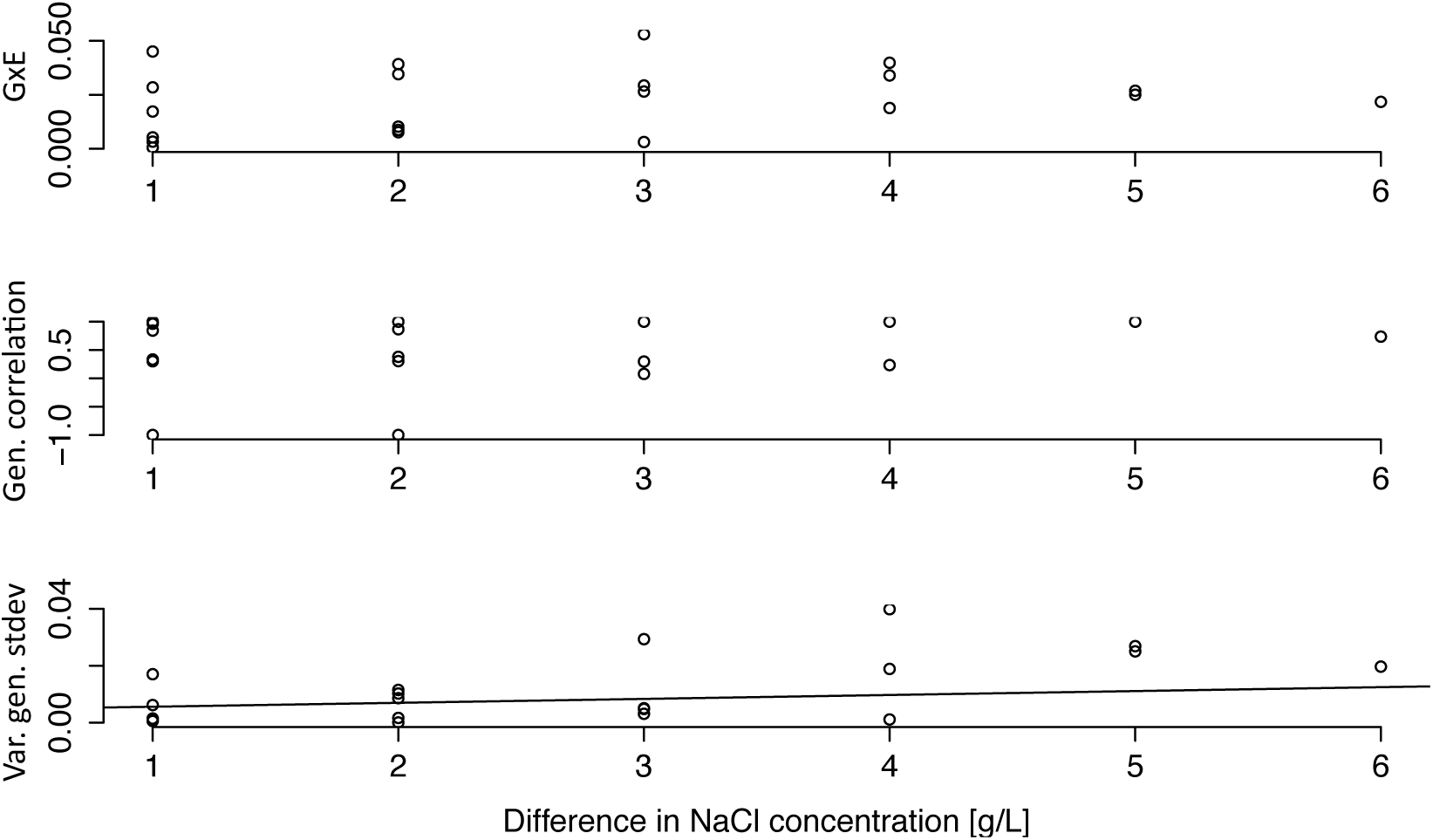
Overall GxE interactions, genetic correlation of growth and variation of the genetic standard deviation based on the relative fitness of MA genotypes derived from ancestor strain CC-2344 plotted against the differences in NaCl concentrations of the pairs of test environments. Lines indicate a significant relationship between the variables after permutation tests.

### 4. Does genetic variance for relative fitness depend on the environment?

Whilst an examination of GxE interactions provides some information about changes in genetic variance across environments, we can also calculate the level of genetic variance among MA genotypes within each environment directly. Regardless of stressor and the ancestor of the MA genotypes, we did not observe any systematic change in the level of expressed genetic variation as environmental stress increased (Fig. 4, linear mixed model, effect of stress level onto expressed genetic variation: *p* = 0.36, effect of stress level onto the variation of the expressed genetic variation: *p* = 0.35). Likewise, there was no significant impact of stress level on evolvability (the variance scaled by the mean, *p* = 0.85).

**Figure 4:**
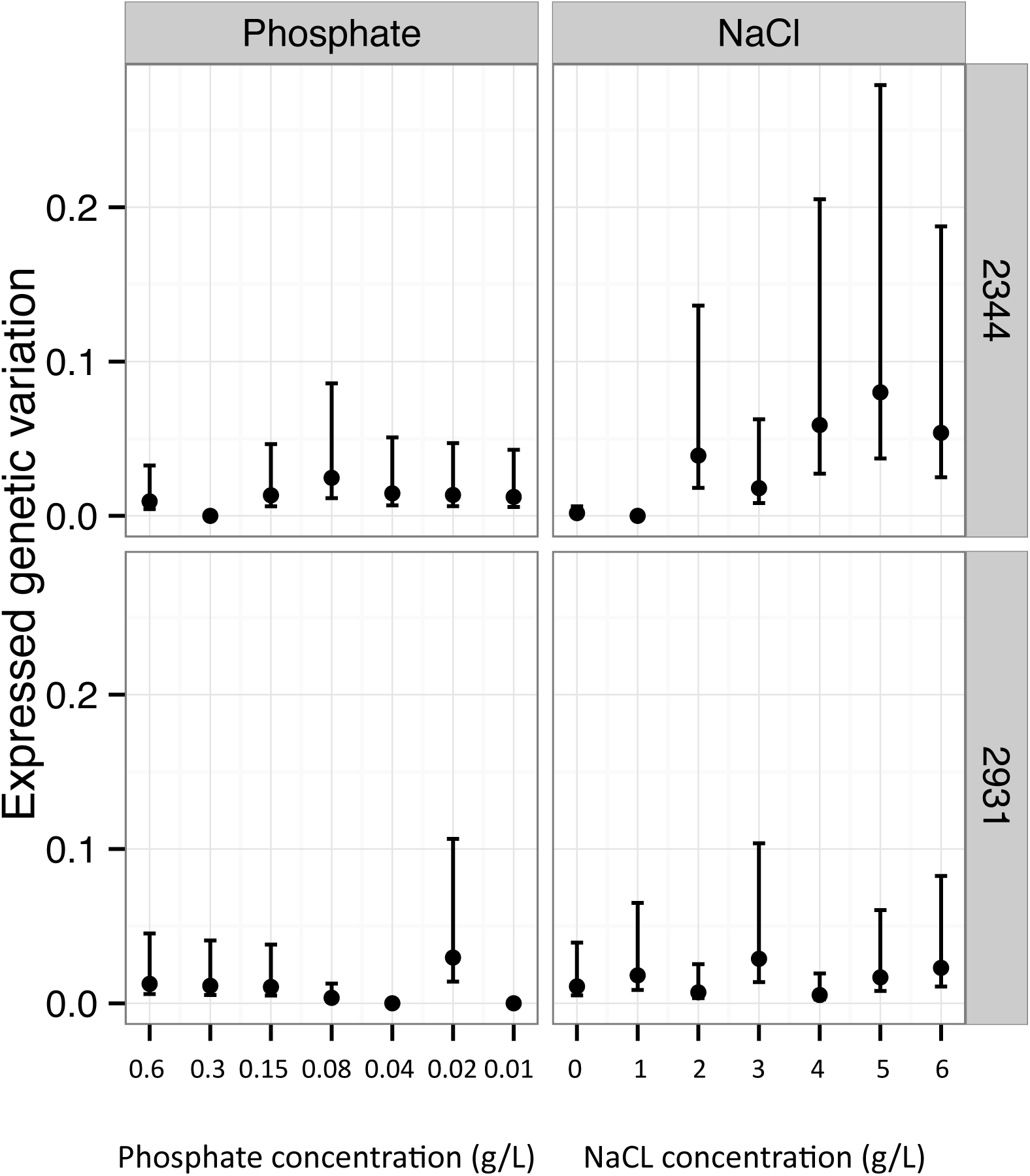
Mean expressed genetic variation of all MA genotypes under different levels of NaCl or phosphate stress, ordered by increase in stress level. Error bars indicate 95% confidence intervals.

## Discussion

Assessing the effect of mutations on fitness across gradients of environmental stress provides a more complete picture of the consequences of new mutations relevant to natural populations than an experiment done in a single environment. Here, we assayed fitness responses of MA lines carrying significant mutational load (Ness et al., 2012; Morgan et al., 2014) across two distinct stress gradients, increasing NaCl concentration and decreasing phosphate concentration. In agreement with previous work (Bell, 1992; Goho and Bell, 2000; Moser and Bell, 2011), both gradients significantly and substantially reduce the fitness of both the MA lines and their ancestors.

Previous studies of mutational effects under deteriorating environmental conditions have often assessed fitness based either directly on maximum growth rates (Fudala and Korona, 2009; Kishony and Leibler, 2003; Szafraniec et al., 2001) or on Malthusian growth parameters per unit of time (Cooper et al., 2005; Jasnos et al., 2008; Remold and Lenski, 2001). However, such time-based measures of fitness do not take into account the likely impact of stressful conditions on generation time (Chevin, 2011). Thus, to avoid overestimation of per-generation selection strength, relative fitness should be scaled by the ancestral generation time under identical conditions.

The results of our study underscore the importance of this distinction. Basing our analyses on growth rates alone, we found no evidence that *de novo* mutations changed growth rates across gradients of the two stressors, as evident by a lack of a significant interaction between the decrease in growth rate and whether or not genotypes had accumulated mutations. Similarly, analyses based on Malthusian fitness parameters showed no effects of stress level treatments on relative fitness. However, relative fitness corrected by ancestral generation times under the same conditions was impacted by environmental deterioration along one of the two stress gradients we investigated. In other words, the increase in generation time caused by stress means that the relatively uniform effects of mutations on growth rates seen across the gradient translates into a greater reduction in absolute fitness per generation since generation time is longer in the more stressful environments. This in turn implies that we would see a greater change of gene frequencies between generations in stressful environments. Specifically, we observed a significant decrease in relative fitness in the highest NaCl concentration investigated. This environment causes an on average 84% decrease in growth rate and is close to the lethal limit for most MA lines. While there is some suggestion that high phosphate stress causes a decrease in relative fitness in MA line genotypes derived from one of the two ancestors, this effect was not significant. The highest phosphate stress caused a smaller fitness decrease in the ancestors (65%) than our high NaCl stress (84%), so this difference may simply reflect a difference in the severity of stress that our gradients imposed. A systematic comparison of mutational effects and severity of the stressful environment remains to be undertaken to disentangle these factors. Overall, these results suggest that changing environmental conditions can exacerbate mutational effects, but that this effect is not necessarily linear and only detectable under near-lethal circumstance. Moreover, in a broader sense, this result may help to explain the inconsistency of results obtained in some previous studies, which only investigated mutational effects under one control and one stressful environment (Baer et al., 2006; Kishony and Leibler, 2003; Remold and Lenski, 2001).

We also tested whether increased environmental stress affected the expressed genetic variation both between and within environments. Only MA genotypes derived from one of the two ancestors showed significant GxE interactions, and only under NaCl stress. When we partitioned these interactions into their components, they seemed to be mainly driven by an increase of the variation of the genetic standard deviation of genotype fitness across increasingly different NaCl concentrations. However, all other MA genotypes and stressors investigated did not show any significant genotype-by-environment interactions. Likewise, we saw no systematic change of the expressed genetic variation within environments across both stress gradients. Both these findings are consistent with previous results (Latta et al., 2015), in which only a minority of test strains and environments yielded significant GxE interactions. Our results are in part consistent with theoretical expectations formulated by Martin and Lenormand (Martin and Lenormand, 2006), who found that stress does not impact the average mutational effect (though it might very well interact with the effects of individual mutations). In contrast to their findings we do not detect a significant effect of stress on the expressed genetic variation. It is noteworthy, however, that lines and stress conditions that had the strongest effect on relative fitness in our experiment also show an increase of expressed genetic variation under deteriorating conditions. Thus, changes in expressed genetic variation under environmental deterioration might be both depending on the genetic background as well as the exact nature of the stress. We find our results of limited effects of non-density dependent stressors on selection to be in accordance with expectations formulated by Agrawal and Whitlock (2010). However, we did not contrast density and non-density dependent stressors explicitly here.

Although mutations mostly had a constant effect across stress levels, fitness was affected both by the type of stress and the ancestor from which a MA genotype was derived. Thus, accurately predicting fitness responses based on the number of mutations might require a mechanistic understanding of the gene in which a given mutation resides. For example, mutational effects might be amplified if the gene carrying them is part of the core genome, whereas the effects of mutations in condition-dependent genes will be harder to detect, and depend on the assay conditions. Likewise, effects of mutations in highly connected genes may be impacted by the gene’s epistatic interactions (Draghi and Whitlock, 2015). Moreover, the effect of a mutation might depend on the genetic background in which the mutation occurs. For example, if the ancestral genotype were close to a fitness peak in its native environment, most mutations are expected to be deleterious (Fisher, 1958), therefore, measuring fitness in a novel stressful environment where the ancestor is not well adapted will alter our interpretation of a mutation’s effect on fitness. Lastly, different stresses will impact the physiological state of the cell differently and thus might have different interactions with the mutations present.

In summary, these results suggest that new mutations obtained by mutation accumulation may have environment-specific effects on relative fitness. The implications of this are two fold. First, a stress-mediated release of cryptic genetic variation is possible, especially under highly detrimental conditions. Such a release may, on the one hand, increase adaptiveness of a population under stress by increasing variation between individuals. On the other hand, however, it may represent a genetic load that might decrease the fitness of a stressed population below a recovery threshold. Secondly, interactions between genotypes and test environments were both ancestor- and stress-specific, thus making generalizations about the adaptive nature of GxE interactions difficult. The environments we used have shown GxE interaction amongst natural isolates of *C. reinhardtii* (Bell, 1992; Goho and Bell, 2000; Moser and Bell, 2011) but, overall, we found GxE interactions in our study to be of limited importance. This provides a hint at the importance of selection in maintaining GxE in natural populations: mutations responsible for much of the GxE interactions seen in genotypes isolated from nature seem to be actively maintained by selection, rather than due to the accumulation of conditionally deleterious mutations that are neutral in the environment that they arise in (except for highly detrimental conditions). More broadly, it implies that our understanding of the distribution of mutational effects gained from laboratory studies (Kassen and Bataillon, 2006; Latta et al., 2015; Sanjuan et al., 2004; Wloch et al., 2001; Zeyl and DeVisser, 2001) may be representative of the situation in more natural conditions.

